# The common neoantigens in colorectal cancer are predicted and validated to be presented or immunogenic

**DOI:** 10.1101/682617

**Authors:** Zhaoduan Liang, Lili Qin, Lei Chen, Wenhui Li, Chao Chen, Yaling Huang, Le Zhang, Songming Liu, Si Qiu, Yuping Ge, Wenting Peng, Xinxin Lin, Xuan Dong, Xiuqing Zhang, Bo Li

## Abstract

Colorectal cancer (CRC) is a malignant cancer with high incidence and mortality in the world, as the result of the traditional treatments. Immunotherapy targeting neoantigens can induce durable tumor regression in cancer patients, but is almost limited to individual treatment, resulting from the unique neoantigens. Many shared oncogenic mutations are detected, but whether the common neoantigens can be identified in CRC is unknown. Using the somatic mutations data from 321 CRC patients combined with a filter standard and 7 predicted algorithms, we screened and obtained 25 HLA-A*11:01 restricted common neoantigens with high binding affinity (IC50<50 nM) and presentation score (>0.9). Except the positive epitope KRAS_G12V_8-16_, 11 out of 25 common neoantigens were proved to be naturally processed and presented on constructed K562 cell surface by mass spectroscopy (MS), and 11 out of 25 common neoantigens specifically induced *in vitro* pre-stimulated cytotoxic lymphocyte (CTL) to secrete IFN-γ. However, only 2 out of 25 common neoantigens were simultaneously presented and immunogenic. Moreover, using cell-sorting technology combined with single-cell RNA sequencing, the immune repertoire profiles of C1orf170_S418G_413-421_ and KRAS_G12V_8-16_-specific CTL were clarified. Therefore, common neoantigens with presentation and immunogenicity could be found in CRC, which would be developed as the universal targets for CRC immunotherapy.

## Introduction

Colorectal cancer (CRC) is the third commonest diagnosed malignant cancer and the second leading cause of cancer death in the world [1]. In 2018, more than 1.8 million new cases of CRC and almost 881 thousand cases of CRC-interrelated death occurred in the world [1], and the global burden of CRC is estimated to reach over 2.2 million new cases and 1.1 million cancer deaths by 2030 [2]. Traditionally surgical resection can cure the early stage of CRC, but about 50% of patients ultimately die of distant metastases. While chemotherapy, radiation therapy and targeted therapy can extend overall survival, less than 15% of patients with metastatic CRC survive beyond 5 years [3]. Therefore, the novel and more effective therapeutic approaches for CRC are necessary to develop.

In the recent years, based on a better knowledge of the complex interactions between the immune system and the tumor microenvironment, immunotherapy has become a novel effective and promising therapeutic strategy for cancer, and its efficacy is widely tested by CRC model. The vast majority of CRC patients with deficient mismatch repair (dMMR) or highly microsatellite instable (MSI-H) benefit from immune checkpoint inhibitors, which is not effective in other CRC patients with proficient MMR (pMMR) or microsatellite stable (MSS) [4]. Patients with CRC do not respond to autologous tumor lysate DC (ADC) and peptide vaccines [4]. T cells, which are engineered to express an affinity-enhanced T-cell receptor (TCR) or an antibody-based chimeric antigen receptor (CAR) targeting tumor associated antigens (TAAs), such as carcinoembryonic antigen (CEA) [5, 6] and human epidermal growth factor receptor-2 (HER2) [7], regress metastatic CRC, but simultaneously mediate severe autoimmunity in patients. These results highlight the importance of identifying tumor specific antigens, which optimally discriminate tumor and normal tissues, for improving the efficacy and safety of adoptive T-cell therapy for CRC.

Compared with TAAs, which lowly express in some normal cells but overexpress in tumor cells [5–7], mutated tumor-specific antigens (TSAs) arise from the somatic mutations in protein-coding regions of tumor cells, and are exclusively present in malignant cells and not produced by normal tissues [8]. The accumulated mutations in cancers include nonsynonymous, nonsense, indel and frame shift, and are classified into driver mutations, which involve in uncontrolled cell growth and tumor metastasis, and passenger mutations, which may not contribute to the tumorigenic phenotype, but increase immunogenicity [9]. Through the antigen presentation system, peptides, which contain the mutant sites and are called neoantigens, can be presented on the surface of tumor cells by major histocompatibility complex (MHC) molecules, then recognized by T cells to lead robust anti-metastatic CRC activity [10]. Furthermore, T-cell responses elicited by neoantigens are not subject to host central tolerance in the thymus and also bring fewer toxicities deriving from autoimmune reactions to normal cells [11]. The highly individual neoantigens actualize the personalized cancer immunotherapies, but limited the development of “one fits all” pharmacologic solutions [11].

It has been shown that some cancers with high tumor mutational burden (TMB) possess a set of common neoantigens owing to the microsatellites [11]. The presence of microsatellite instability has been found in approximately 15-20% CRC, and dMMR CRC has a high TMB, which is far higher than the standard value [4]. Furthermore, driver mutations possess only 8% CD8^+^T-cell neo-epitopes, while passenger mutations possess 92% CD8^+^ T-cell neo-epitopes and 100% CD4^+^T-cell neo-epitopes [9]. HLA-A*11:01 allele has high prevalence in US Caucasians, Asian-Americans and China [12] (http://www.allelefrequencies.net/). Therefore, we hypothesized that targeting the common neoantigens, comprising driver mutations and passenger mutations and restricted by HLA-A*11:01, not only improved the efficacy, safety and adoption of CRC immunotherapy, but also reduced the cost and time of CRC clinical treatment. In the present study, we predicted the HLA-A*11:01 restricted common neoantigens from the somatic mutations data of 321 patients with CRC and validated their presentation and immunogenicity, which would become the new targets for CRC immunotherapy.

## Materials and methods

### Cell lines

The TAP-deficient T2 cell line (CRL-1992), K562 cell line (CCL-243) and HEK-293 cell line (CRL-1573) were purchased from the American Type Culture Collection (ATCC), and respectively maintained in Iscove’s Modified Dulbecco’s Medium (IMDM, Gibco), RPMI-1640 medium (Gibco) and Dulbecco’s Modified Eagle’s Medium (DMEM, Gibco) with 10% fetal bovine serum (FBS; Hyclone) at 37°C in a humidified 5% CO_2_ incubator. T2 cell line and K562 cell line were retrovirally transduced with retrovirus encoding HLA-A*11:01. Cells were authenticated by HLA genotyping, tested for mycoplasma by PCR method, and maintained in medium no more than 2 months from each thaw. Human peripheral blood was obtained from anonymous healthy donors who had signed informed consents. Peripheral blood mononuclear cells (PBMCs) were isolated by Ficoll-Hypaque gradient centrifugation and maintained in RPMI-1640 medium supplemented with 10% FBS at 37 °C in a humidified 5% CO_2_ incubator. The study was approved and conducted by Institutional Review Board of Beijing Genomics of Institute (BGI)-Shenzhen (No. BGI-IRB18142).

### Mutation selection and epitope prediction

The somatic mutations data of 321 patients with CRC from China-Colorectal Cancer Project (COCA-CN, https://icgc.org/icgc/cgp/73/371/1001733) in ICGC (International Cancer Genome Consortium) database (http://icgc.org) were downloaded and further analyzed. In briefly, missense variants that caused amino acid changes in coding regions were filtered according to a standard, in which the frequency of single-nucleotide variants (SNVs) was over 5 out of 321 patients and insertions or deletions (InDels) was over 2 out of 321 patients. After obtaining the list of the tumor-specific mutant proteins, we extracted the peptide sequences around the mutated sites. As MHC class I molecules bind to peptides 9-10 amino acids in length with the highest affinity [13], peptides were extracted *in silico* from 19 amino acids sequences, with 9 amino acids upstream and 9 amino acids downstream of mutated amino acids, and 19 sets of 9-mer or 10-mer sequence containing mutated site from each mutated protein were identified and predicted by algorithms. Wild-type peptides with the same length as mutated peptides were extracted as references. The potential binding affinity between extracted peptides and HLA-A*11:01 allele was analyzed simultaneously by NetMHC-4.0 [14], NetMHCpan-3.0 [15], NetMHCpan-4.0 [16], PSSMHCpan-1.0 [13], PickPocket-1.0 [17] and SMM [18]. Results were exhibited as predicted equilibrium binding constants of IC50 (50% inhibitory concentration, nM), in which strong binders meant the predicted binding affinity IC50 values were less than 50 nM, and weak binders were that of 50-500 nM [19]. Moreover, we used our software Epitope Presentation Integrated prediCtion (EPIC) [20], with a fixed expression value 4 Transcripts Per Kilobase Million (TPM) as inputs, to predict the presentation of extracted peptides, in which the results were shown by the scores of highest presentation probability. These mutant peptides with strong binding capacity and high score of presentation probability were selected as potential neoantigens.

### The construction of putative neoantigens transduced K562 cells (HLA-A*11:01^+^)

Six predicted neoantigens were linked into a tandem neoantigen as previously described [21]. Briefly, six predicted neoantigens, which each had 27 amino acids with the mutation at position 14, were connected by a start linker (GGSGGGGSGG), middle linkers (GGSGGGGSGG) and an end linker (GGSLGGGGSG). The N terminal of a tandem neoantigen was successively linked with the kozak sequence (GCCACC) and the signal peptide sequence (SPMRVTAPRTLILLLSGALALTETWAGS), and the C terminal was linked with the MHC class I trafficking signal (MITD) sequence (IVGIVAGLAVLAVVVIGAVVATVMCRRKSSGGKGGSYSQAASSDSAQGSDVSLTA) and termination codon. The tandem minigene DNA fragment coding the above sequence was synthetized and cloned into lentiviral expressing vector pLVX (CMV-EF1a-ZsGreen-P2A-Bsd; provided by Viraltherapy Technologies Ltd, Wuhan, China). Lentiviral particles encoding tandem minigenes were produced from HEK-293 cells, which were simultaneously transduced with packaging constructs (RRE, REV and VSVG (invitrogen)) and the expressing vector, and infected mono HLA-A*11:01 allelic K562 cells. Positive K562 cells were selected by Blasticidine S hydrochloride (5 μg/ml, Sigma) and detected through the percentage of reporter gene ZsGreen.

### The preparation of MHC Class I bound peptides

MHC-I peptidomes were obtained from mono HLA-A*11:01 allelic K562 cells transduced with putative neoantigens as described previously [22]. In brief, 1×10^9^ cells were dissociated using lysis buffer (0.25% sodium deoxycholate, 1% n-octyl glucoside, 100 mM PMSF and protease inhibitors cocktail in phosphate buffer saline (PBS)) at 4 °C for 60 min. Lysate were further cleared by centrifugation at 14,000 g for 30 min. Cleared lysate were purified with anti-pan-HLA class I complexes antibody (clone W6/32), which was covalently bound Protein-A Sepharose CL-4B beads (GE Healthcare). Beads were washed with Tris-HCl buffer containing NaCl. The MHC-I molecules were eluted at room temperature using 0.1 N acetic acid. Eluate was loaded on Sep-Pak tC18 cartridges (Waters, 100 mg). The C18 cartridges were first washed with 0.1% TFA, then with 0.1% TFA containing 30% ACN to separate peptides from MHC-I complexes. Eluate was concentrated to 20 μl using vacuum centrifugation. Finally, 5 μl of sample was used for Parallel Reaction Monitoring (PRM) mass analysis.

### Peptide validation by mass spectroscopy (MS) analysis with PRM

Peptides were separated by a nanoflow HPLC (15 cm long, 75 μm inner diameter column with ReproSil-Pur C18-AQ 1.9 μm resin) and coupled on-line to a Fusion Lumos mass spectrometer (Proxeon Biosystems, Thermo Fisher Scientific) with a nanoelectrospray ion source (Proxeon Biosystems). Peptides were eluted with a linear gradient of 5-80% buffer B (98% ACN and 0.1% FA) at a flow rate of 500 nl/min over 3 hours. Data of each injection was acquired using a corresponding transition list (data not shown). Full scan MS spectra were acquired at a resolution of 6,000 at 350-1,400 m/z with a target value of 4×10^5^ ions. MS/MS resolution was 60,000 at 150-2,000 m/z, and higher collisional dissociation (HCD) was employed for ion fragmentation. The interpretation of MS data was performed with Skyline. To validate a peptide which could be presented by MHC-I complex, the following criteria were considered: i) the variation of retention time between precursor ions was less than 3 min; ii) the pattern and retention time were matched between synthetic peptide and target peptide for no less than 5 product ions.

### The preparation of tetramer of peptide-MHC complex

Peptides (Table 1) and HLA-A*11:01-restricted KRAS G12V_8-16_ (VVGAVGVGK) as positive peptide were synthesized from GenScript (Nanjing, China), with purity greater than 98% by mass spectroscopy. Peptide-MHC tetramers were generated as previously described [23]. In briefly, Peptides (400 μM) were mixed with Flex-T™ HLA-A*11:01 Monomer UVX (Biolegend), then subjected to UV light for 30 min on ice. The MHC monomers exchanged with peptides were tetramerized in the presence of allophycocyanin (APC) conjugated streptavidin (BD Biosciences) for 30 min at 37 °C, then the reaction was stopped by PBS containing D-Biotin (500 μM) and NaN_3_ (10%), then kept at 4 °C overnight for use.

**Table 1.**
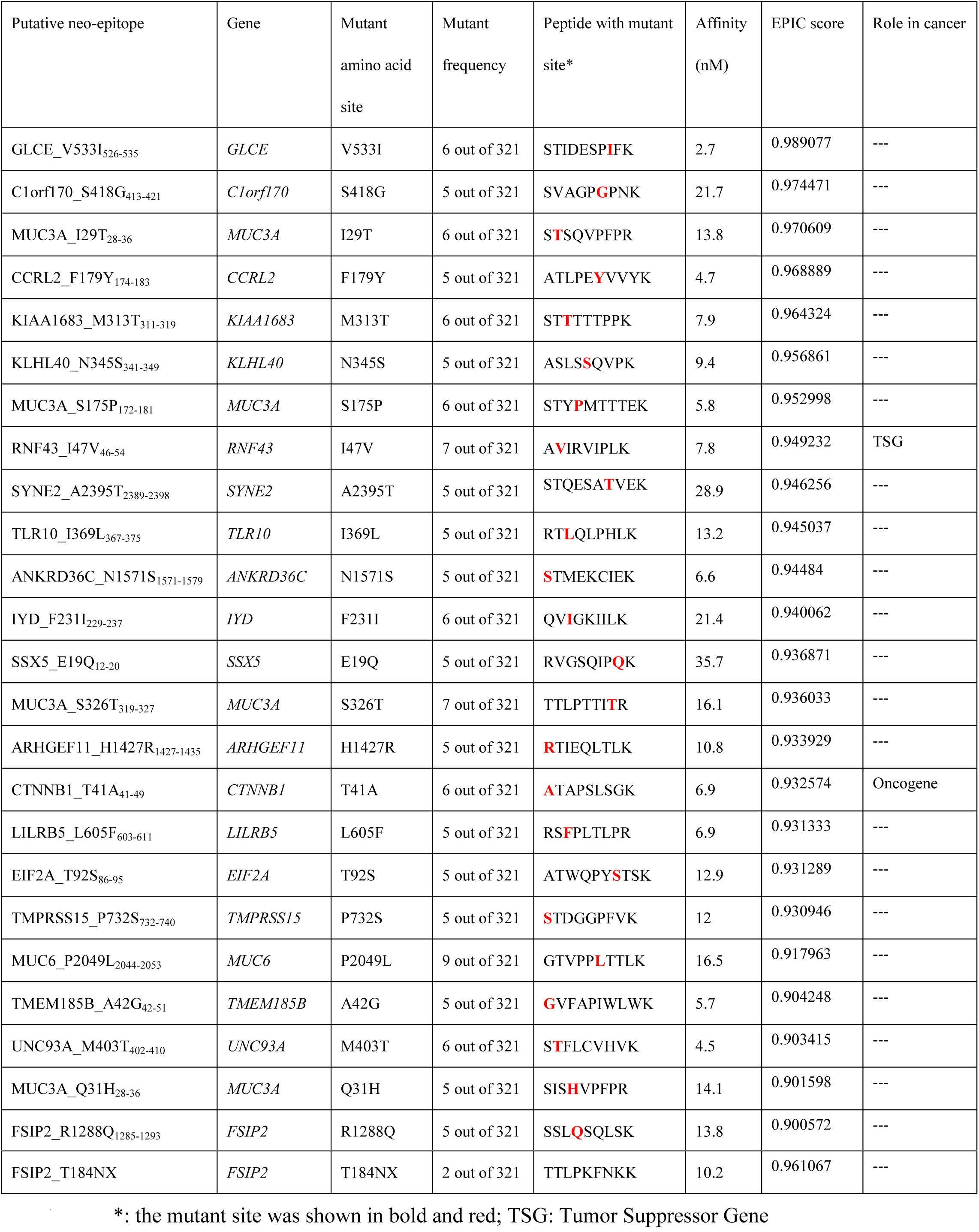
The list of the common predicted peptide candidates of CRC

### The generation of mature dendritic cells (mDCs)

Monocytes (CD14 positive) were positively selected using CD14 MicroBeads (Miltenyi Biotec) from PBMCs of healthy donors as the manufacturer’s protocol, and cultured in CellGenix^TM^ DC media (CellGenix) supplemented with 2% human serum albumin (HSA, CSL Behring L.L.C.), granulocyte-macrophage colony stimulating factor (GM-CSF, 100 ng/ml; PeproTech) and interleukin 4 (IL-4, 100 ng/ml; PeproTech) for 5 days. On day 6, immature DCs were stimulated to mature by TNF-α (10 ng/ml; PeproTech), IL-6 (50 ng/ml; PeproTech), IL-1β (10 ng/ml; PeproTech), Prostaglandin E2 (500 ng/ml; Sigma) and Poly(I:C) (10 μg/ml; InvivoGen) for 2 days. After harvest, mDCs were pulsed with peptides (1 μg/ml, Table 1 and KRAS G12V_8-16_) in FBS-free RPMI-1640 medium for 4 hours at 37 °C, and used as the antigen presented cells. The maturity of DCs was determined through the morphology and the phenotype of the expressions of CD80, CD83, CD86, CD11c and HLA-DR.

### The induction of neoantigen-specific cytotoxic lymphocyte (CTL)

CD8^+^T cells were positively enriched using CD8 MicroBeads (Miltenyi Biotec) from PBMCs of healthy donors as the manufacturer’s protocol, stimulated by mDCs pre-loaded with peptides at a 4:1 ratio, and maintained in HIPP™-T009 medium (Bioengine) in the presence of 2% autoserum and IL-21 (30 ng/ml, PeproTech) in a 37 °C 5% CO_2_ incubator for 12 days. On day3, the co-culture system was supplemented with IL-2 (5 ng/ml, PeproTech), IL-7 (10 ng/ml, PeproTech) and IL-15 (10 ng/ml, PeproTech), which were repeated every 2-3 days. After 12-day culture, the pre-stimulated CD8^+^T cells were re-stimulated with same peptide-pulsed mDCs as above and incubated for another 12 days.

### Enzyme-linked immunospot (ELISPOT) assay for IFN-γ

IFN-γ ELISPOT assay strip plate which was pre-coated with anti-human IFN-γ mAb (1-D1K, Mabtech) was washed with PBS and blocked with RPMI-1640 containing 10% FBS. CTLs were co-cultured with T2 cells pre-pulsed with or without peptides (10 μg/ml) in the above pre-treated ELISPOT plate for 24 hours. Each sample was set with repetition. Plate was rinsed with PBS, then added with alkaline phosphatase (ALP) labeled anti-human IFN-γ mAb (7-B6-1-ALP, 1:200; Mabtech) for 2 hours. After rinsing, 5-Bromo-4-chloro-3-indolyl phosphate/Nitro blue tetrazolium (BCIP/NBT, Mabtech) was used to develop the immune-spot according to the manufacturer’s protocol. Spots were imaged and counted by an ELISPOT Reader (BioReader 4000, BIOSYS). Positive response was judged according to that the number of specific spots was more than 10 and at least two-fold greater than that of negative control [24].

### Fluorescence-activated cell sorting (FACS)

Cells were collected, washed and resuspended in PBS containing 2% FBS (FACS buffer). Cells were stained with fluorescent dye conjugated antibodies for 15 min at 4 °C. PE conjugated anti-CD8 antibody, APC conjugated pMHC tetramer, APC conjugated anti-CD86 antibody, PE conjugated anti-CD83 antibody, PE conjugated anti-CD80 antibody, APC conjugated anti-CD11c antibody, PE conjugated anti HLA-DR antibody, and isotype matched antibodies were used in this study and purchased from BD Biosciences. After washing twice in FACS buffer, Cells were analyzed using a FACSAria II(BD Biosciences) with live cell gating based on 4’,6-diamidino-2-phenylindole (DAPI) exclusion, and CD8^+^pMHC tetramer^+^ cells were sorted for single-cell RNA sequencing. The data were analyzed using FlowJo software (Tree Star).

### The analysis of neoantigen-specific T-cell receptor repertoire by single-cell RNA sequencing

According to the manufacturer’s protocol of Chromium™ Single Cell V(D)J Reagent Kits (10x Genomics, Inc.), sorted CD8^+^pMHC tetramer^+^ T cells were partitioned and captured into the Gel Bead in Emulsion (GEM) through the rapid and efficient microfluidics technology of the Chromium™ single-cell controller (10x Genomics, Inc.). Single cell and the Gel Bead were lysed in the GEM, then the contents of the GEM were incubated in the Reverse Transcription-Polymerase Chain Reaction (RT-PCR) to generate full-length, and mRNA transcripts were barcoded on their poly A-tails. Barcoded cDNA molecules were pooled after GEMs being broken, and full-length V(D)J segments from TCR cDNA were enriched by PCR amplification and constructed as a library for Illumina^®^-ready sequencing. TCR repertoire and paired TCR were analyzed by the Cell Ranger™ analysis pipelines. The gene usage was assigned using the IMGT nomenclature.

## Results

### The selection of mutant candidate peptides

In order to analyze the potential common neoantigens of CRC in China, which derived from driver mutations or passenger mutations, we ultimately collected 3,500 SNVs, of which the frequency was over 5 out of 321 patients, and 191 InDels, of which the frequency was over 2 out of 321 patients, from the somatic mutation data of 321 CRC patients from COCA-CN in ICGC database on August 29, 2018. Both 9-10-mer mutant epitope candidates and reference peptides were extracted from 19-amino acid length, with 9 amino acids upstream and 9 amino acids downstream of the 3,691 mutant sites. A total of 60,169 epitopes of SNV and 6,891 epitopes of Indels were generated and predicted the binding affinity with HLA-A*11:01 allele by NetMHC-4.0, NetMHCpan-3.0, NetMHCpan-4.0, PSSMHCpan-1.0, PickPocket-1.0 and SMM simultaneously. As a result, 56 mutant epitopes were selected as the IC50 value of predicted binders was less than 50 nM by at least three software packages, and the smallest affinity predicted value during three softwares was taken as the affinity predicted score of the peptide-MHC complex (Table 1 and data not shown). Furthermore, the algorithm EPIC, which was an effective, flexible and publicly available HLA-I presented epitope prediction method, was used to evaluate the probability of presentation of 56 mutant epitopes with strong binding affinity. The EPIC score value of 25 out of 56 mutant epitopes was more than 0.9, which meant that the probability of predicted epitope being presented by MHC was absolutely high (Table 1). Finally, 25 mutant epitopes related to 25 somatic mutations of 21 genes were selected as peptide candidates, and assessed their presentation and immunogenicity (Table 1). Moreover, during the 21 genes, we found that *RNF43* gene was a tumor suppressor gene, *CTNNB1* gene was an oncogene, and the remaining genes had not been identified as tumor-associated genes, which also encoded potential neoantigens by CRC, in Cancer Gene Census database (https://cancer.sanger.ac.uk/census) (Table 1).

### The expression of predicted peptides from constructed K562 cells

T cells attacking targeted cells mainly depends on that TCR recognize T-cell epitopes, which are expressed, naturally processed and presented by MHC molecules on the cell surface [24]. In order to improve the probability of the predicted epitopes being presented by MHC class I, the predicted peptides were linked into the tandem minigenes, and fused with N-terminal leader peptide and C-terminal MITD, which have been proved to strongly improve the presentation of MHC class I and class II epitopes [25]. The assembled base sequences were cloned into the multiple clone site (MCS) of lentiviral vector (Figure 1(a)), which were operated by CMV promoter and tracked by a reporter gene ZsGreen. As a result, we constructed five tandem-minigene stably transfected K562 cell lines with mono HLA-A*11:01 allele. CRC-1-K562 cells contained the minigenes of GLCE_V533I, C1orf170_S418G, MUC3A_I29T, CCRL2_F179Y, KIAA1683_M313T and KLHL40_N345S; CRC-2-K562 cells contained the minigenes of MUC3A_S175P, RNF43_I47V, SYNE2_A2395T, TLR10_I369L, ANKRD36C_N1571S and IYD_F231I; CRC-3-K562 cells contained the minigenes of SSX5_E19Q, MUC3A_S326T, ARHGEF11_H1427R, CTNNB1_T41A, LILRB5_L605F and EIF2A_T92S; CRC-4-K562 cells contained the minigenes of TMPRSS15_P732S, MUC6_P2049L, TMEM185B_A42G, UNC93A_M403T, MUC3A_Q31H and FSIP2_R1288Q; CRC-5-K562 cells contained the minigenes of TMEM185B_A42G, UNC93A_M403T, MUC3A_Q31H, FSIP2_R1288Q, FSIP2_T184NX and KRAS_G12V (Table 2 and Figure 1(b)). In present study, we selected KRAS_G12V, which is accepted as a common oncogenic mutation and has been proved to be presented by HLA-A*11:01 allele on the basis of T-cell response to KRAS_G12V positive target cells [12], as a positive control of presentation and immunogenicity. In Figure 1(b), the vast majority of tandem-minigene stably transfected K562 cell lines expressed reporter gene ZsGreen, and the range of the expression rate was 85-93%, which was determined by FACS and indirectly reflected the constructed tandem minigenes were expressed in K562 cells.

**Figure 1.**
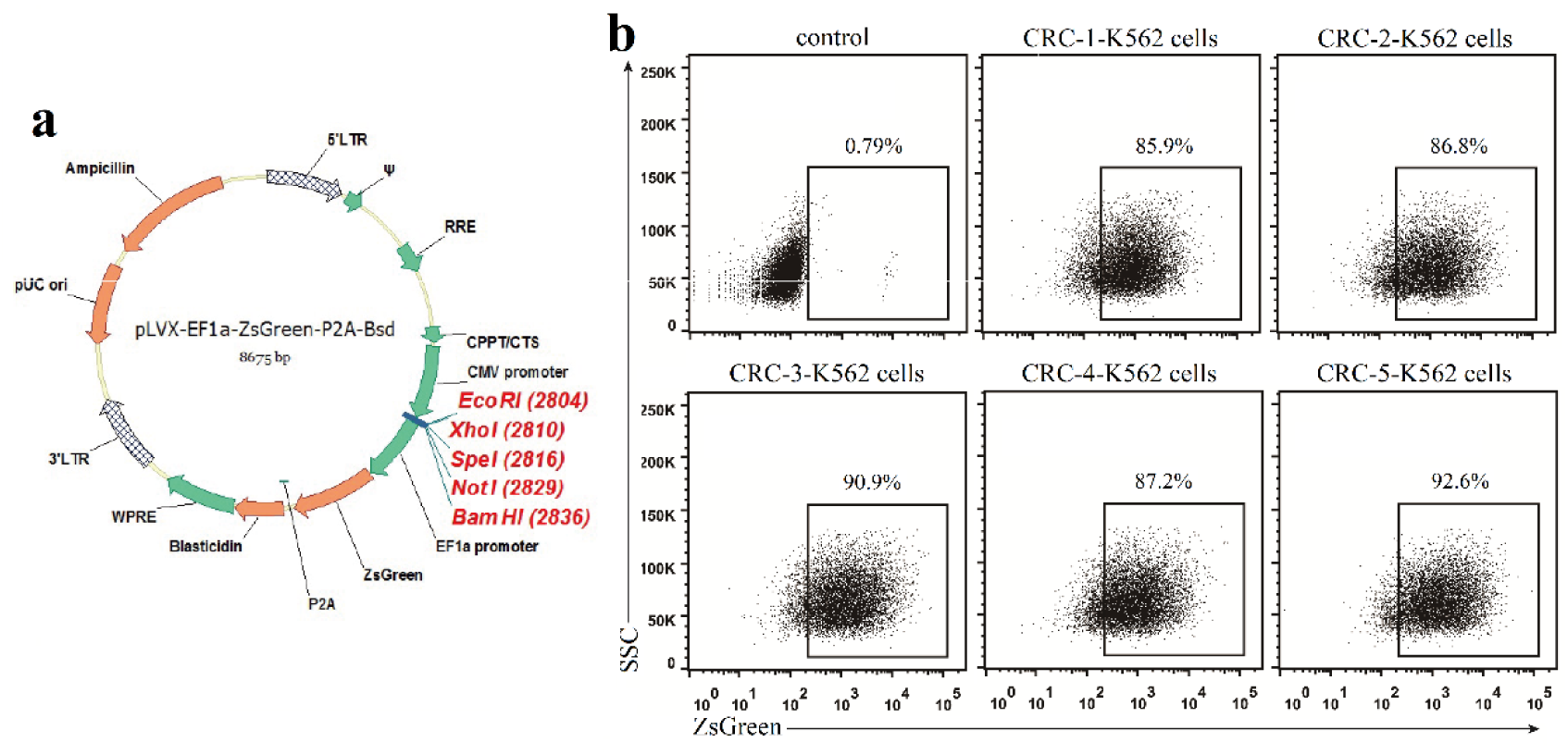
The construction and expression of tandem minigenes encoding predicted neoantigens in K562 cells. (a) The map of the adopted lentiviral vector. The lentiviral vector was gifted and possessed two promoters, of which CMV promoter operated the inserted genes in MCS, and EF1α promoter operated the reporter gene ZsGreen and a screening gene Blasticidine. (b) FACS detected the expression of tandem minigenes in K562 cells. 25 predicted peptides and KRAS_G12V were constructed into five tandem minigenes, packaged into lentivirus and transfected into mono HLA-A*11:01 allelic K562 cells, which resulted in five K562 cell lines, including CRC-1-K562 cells, CRC-2-K562 cells, CRC-3-K562 cells, CRC-4-K562 cells and CRC-5-K562 cells, were obtained (Table 2). The blasticidine-resistance K562 cells expressed ZsGreen, which indirectly reflected the expression of predicted peptides and could be directly tested by FACS.

**Table 2.**
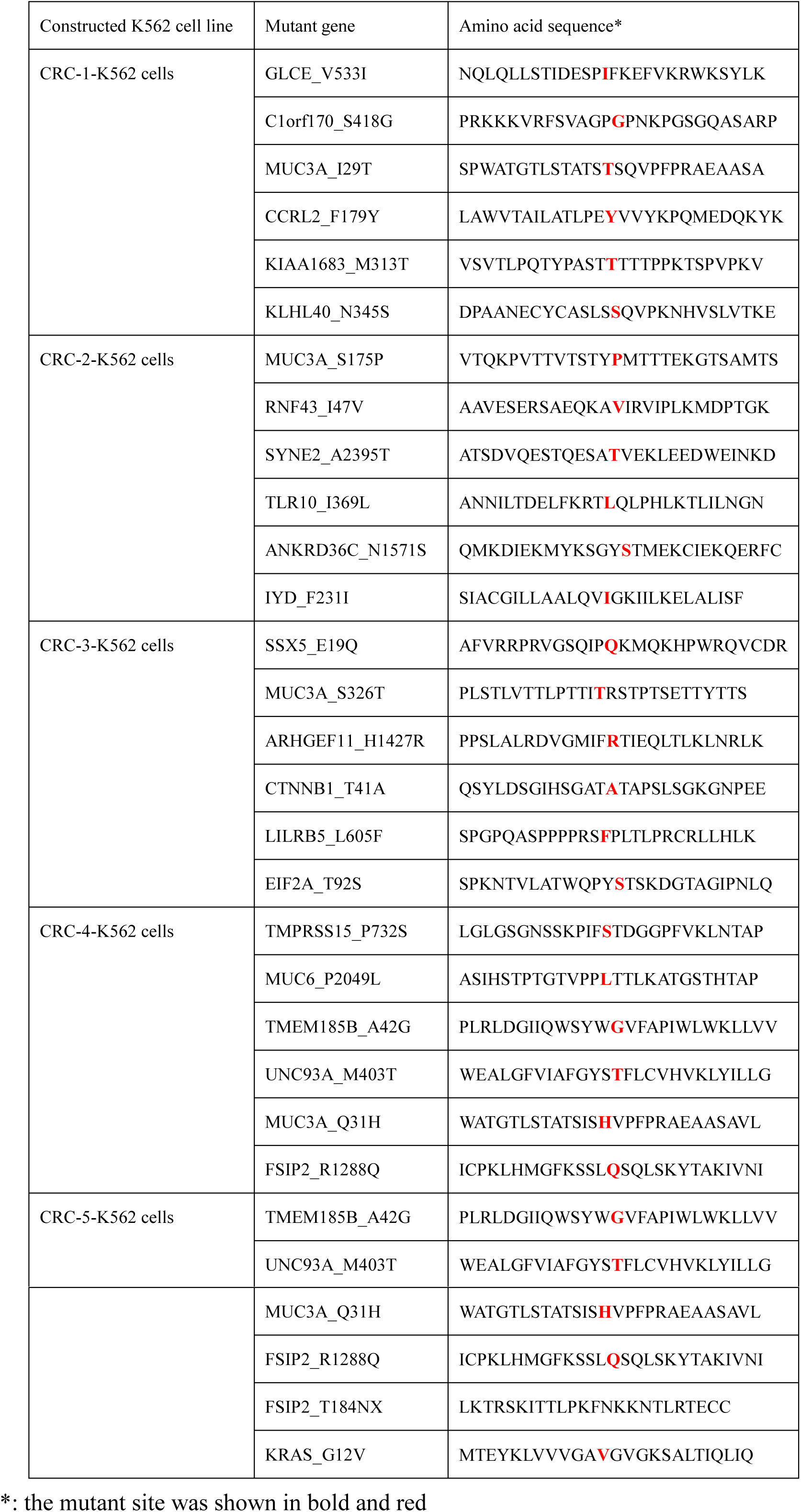
The list of the amino acid sequence of mutant minigenes

### The identification of MHC Class I presented epitopes from constructed K562 cells by MS

In order to verify the predicted epitopes could be naturally processed and presented by cells, we employed an immunoproteomics approach to enrich the immunopeptidome of constructed K562 cells with mono HLA-A*11:01 allele. In this approach, MHC-I restricted peptides were isolated and analyzed with MS. Targeted MS assays with PRM were developed and characterized using stringent search criteria, and resulted in 12 epitopes, including KRAS_G12V_8-16_, GLCE_V533I_526-535_, MUC3A_I29T_28-36_, KLHL40_N345S_341-349_, MUC3A_S175P_172-181_, RNF43_I47V_46-54_, IYD_F231I_229-237_, MUC3A_S326T_319-327_, ARHGEF11_H1427R_1427-1435_, CTNNB1_T41A_41-49_, FSIP2_R1288Q_1285-1293_ and FSIP2_T184NX, were confirmed to be presented by constructed K562 cells and shown as the mirror plot by PDV (Figure 2(a-l))[26].

**Figure 2.**
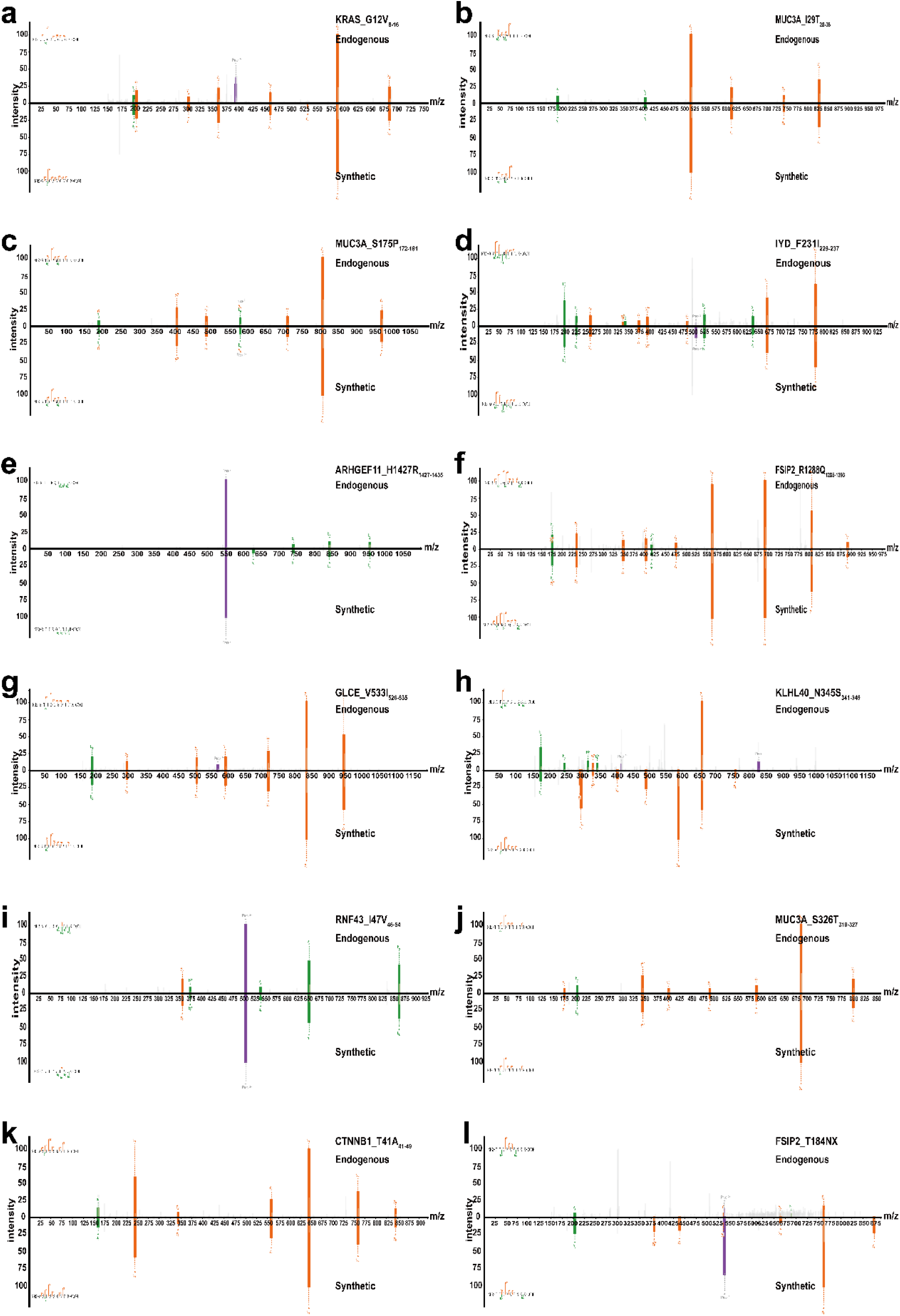
Validation of naturally presented neo-epitopes from constructed K562 cells. The mirror plot of mass spectrometry (MS/MS) chromatographs of the synthetic peptides (bottom) versus theirs experimentally identified analogs (top), including (a) KRAS_G12V_8-16_, (b) MUC3A_I29T_28-36_, (c) MUC3A_S175P_172-181_, (d) IYD_F231I_229-237_, (e) ARHGEF11_H1427R_1427-1435_, (f) FSIP2_R1288Q_1285-1293_, (g) GLCE_V533I_526-535_, (h) KLHL40_N345S_341-349_, (i) RNF43_I47V_46-54_, (j) MUC3A_S326T_319-327_, (k) CTNNB1_T41A_41-49_, and (l) FSIP2_T184NX, were exhibited.

### The analysis of predicted neoantigens activating CTL to secrete IFN-γ in vitro

T cells targeting mutations can be detected from tumor infiltrating lymphocytes (TILs), peripheral memory lymphocytes of cancer patients and peripheral naïve lymphocytes of healthy donors [12, 27]. However, except mouse model, PBMCs from healthy donors were easier obtained to determine the immunogenicity of the predicted neoantigens, comparing to that from patients. To test whether the predicted neoantigens had the characteristic of stimulating CTL to secrete IFN-γ, we used peptide-pulsed mDCs to co-culture with the bulk CD8^+^T cells isolated from HLA-A*11:01^+^ healthy donors for twice in the presence of cytokines, which was supposed to expand neoantigen-reactive CTL. Then CTL were re-stimulated by the corresponding peptide-pulsed T2 cells in the IFN-γ ELISPOT plate. Under inverted phase contrast microscope of 40× object lens, monocytes isolated from PBMCs were observed to display as small round cells, and gradually stretch and adhere the plastic surface of the culture plate on day1(Figure 3(a)). On day8 after the stimulation of a cytokine cocktail for 2 days, mature DCs were exhibited as irregularly large round and suspension cells with blunt, elongate dendritic processes as previously reported (Figure 3(a)) [28, 29]. Furthermore, the phenotype of mature DCs was analyzed by FACS for testing the expression of co-stimulatory molecules and maturation markers. Figure 3(b) showed that more than 98% of DCs expressed CD86, CD80, CD11c and HLA-DR, but a relatively low proportion of mature DC cells expressed CD83 (71.5%), of which the maturation level of DCs was basically reached the international criteria level [30]. After being re-stimulated with T2 cells, which were respectively pre-loaded with epitopes of KRAS_G12V_8-16_, C1orf170_S418G_413-421_, KIAA1683_M313T_311-319_, SSX5_E19Q_12-20_, TMEM185B_A42G_42-51_ and UNC93A_M403T_402-410_, CTL largely secreted IFN-γ, which was manifested by the formation of more than 10 times spots on IFN-γ ELISPOT plate compared with the negative control (Figure 4(a) and (b)). In addition, these epitopes of GLCE_V533I_526-535_, CCRL2_F179Y_174-183_, ANKRD36C_N1571S_1571-1579_, MUC3A_S326T_319-327_, ARHGEF11_H1427R_1427-1435_ and MUC6_P2049L_2044-2053_ also respectively and positively activated CTL to produce low level of IFN-γ, in which the spot number was more than 10 and two-fold greater than that of negative control (Figure 4(a) and (b)). The number of spot mediating by the remainder peptides almost equated that of the negative control, which signified that those peptides did not stimulate CTL to express IFN-γ (Figure 4(a) and (b)). Totally, except the positive control peptide KRAS_G12V_8-16_, 11 out of 25 predicted peptides could activate CTL to secrete IFN-γ *in vitro* and had immunogenicity, which was proved by PBMCs from only one healthy donor, and the positive rate was 44% (11/25).

**Figure 3.**
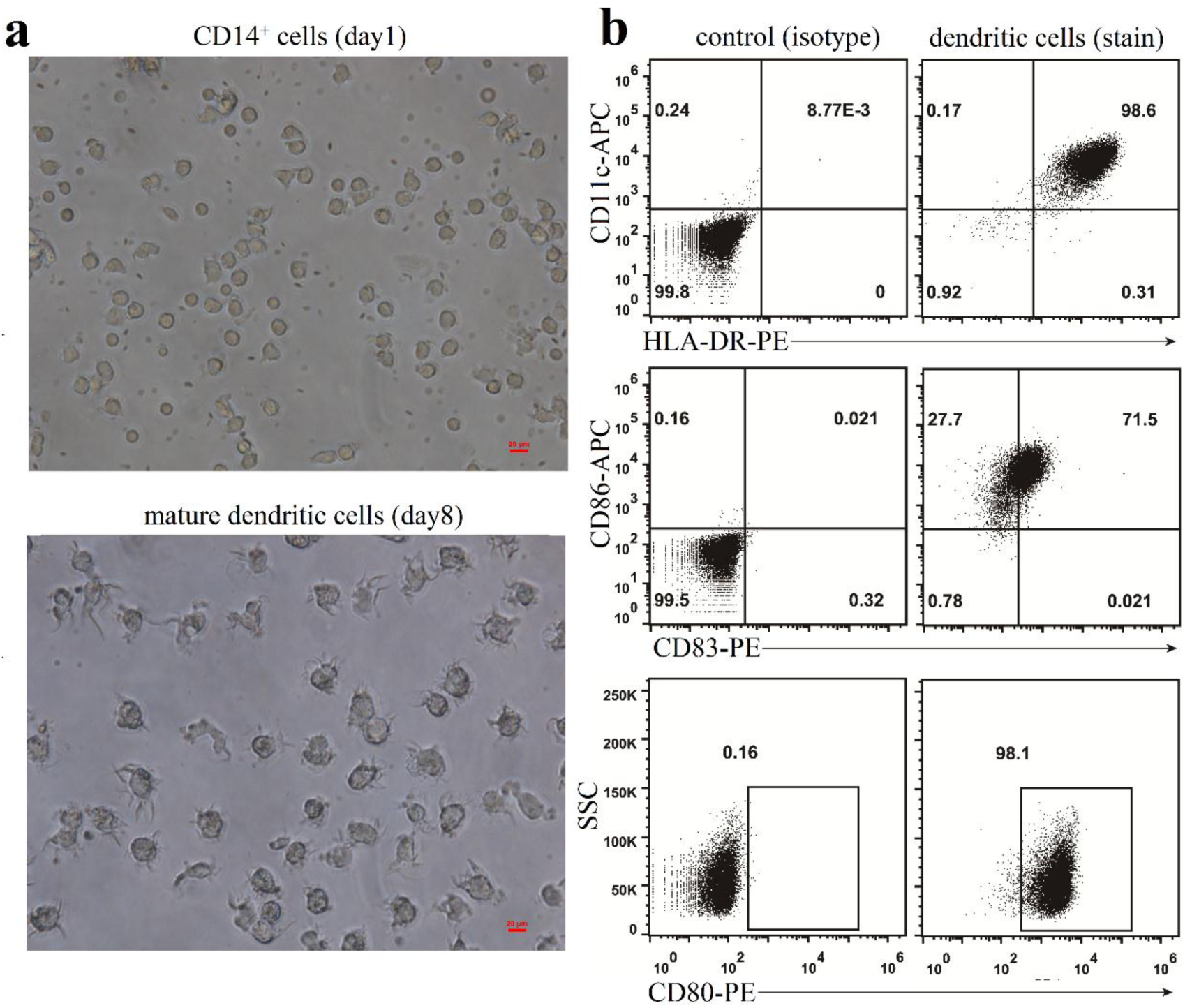
The morphology and the phenotype of mature DCs. (a) The morphology of monocytes and mature dendritic cells. CD14^+^ monocytes on day1 and mature dendritic cells on day8 were observed using inverted phase contrast microscope (Nikon ECLIPSE TS100) and imaged under 40× object lens (scale bar=20 μm). (b) The phenotype of mature dendritic cells. Mature dendritic cells on day8 were collected and stained with fluorescein conjugated antibodies targeting CD11c, HLA-DR, CD86, CD83 and CD80, and the fluorescence signal was detected by FACS.

**Figure 4.**
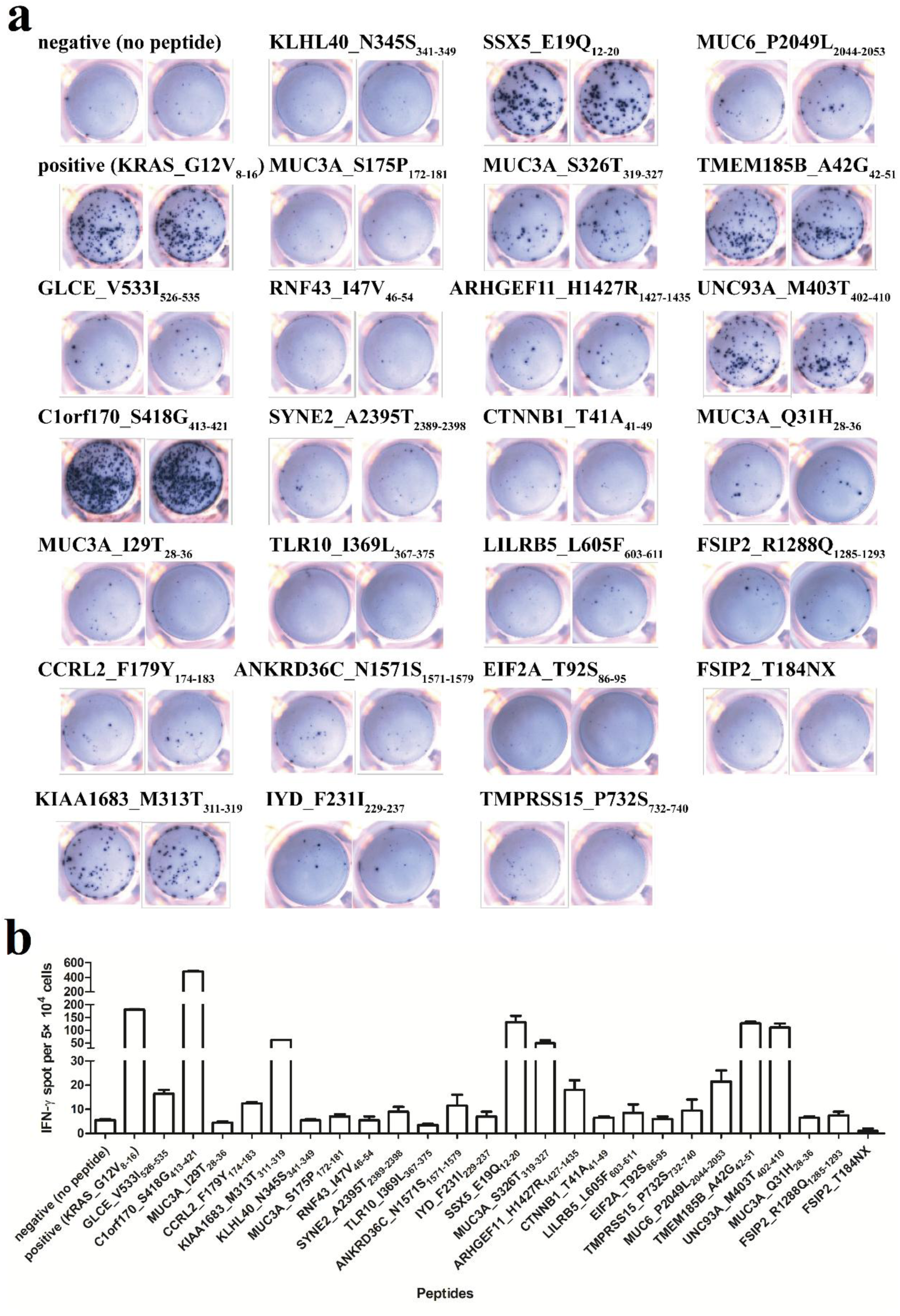
The immune response of CTL to predicted peptide-pulsed T2 cells. (a) The immune-spot diagram of IFN-γ. CTL, which had been co-cultured with peptide-pulsed mDCs, were re-stimulated by peptide-pulsed T2 cells in the IFN-γ ELISPOT plate for 24 hours. The secreted IFN-γ from epitope specific CTL was exhibited as immune spot. There were 25 predicted peptides (Table 1) and a reported positive peptide KRAS_G12V_8-16_ to be tested, and each was in duplicate wells. (b) The statistical number of IFN-γ immune spot from (a). Spots were imaged and counted by an ELISPOT Reader (BioReader 4000, BIOSYS).

### The immune repertoire profiles of epitope-specific CTL from single-cell RNA sequencing

To determine whether epitope-specific CTL were expanded by peptide-pulsed mDCs to the degree of sorting, we selected 6 peptides of KRAS_G12V_8-16_, C1orf170_S418G_413-421_, KIAA1683_M313T_311-319_, SSX5_E19Q_12-20_, TMEM185B_A42G_42-51_ and UNC93A_M403T_402-410_, which strongly activated CTL to secrete IFN-γ (Figure 4(a) and (b)), to prepare tetramer-APC, and stained the bulk CTL with tetramer-APC and CD8-PE simultaneously. We found that the range of the frequency of the epitope-specific CTL after being co-cultured was 0.22-7.08% (Figure 5(a)). Based on the ability of inducing T cells to secrete IFN-γ, we selected to sort KRAS_G12V_8-16_-specific CTL and C1orf170_S418G_413-421_-specific CTL by FACS (Figure 5(b)), of which the sorted cells were single-cell sequenced to determine T-cell receptor repertoires.

**Figure 5.**
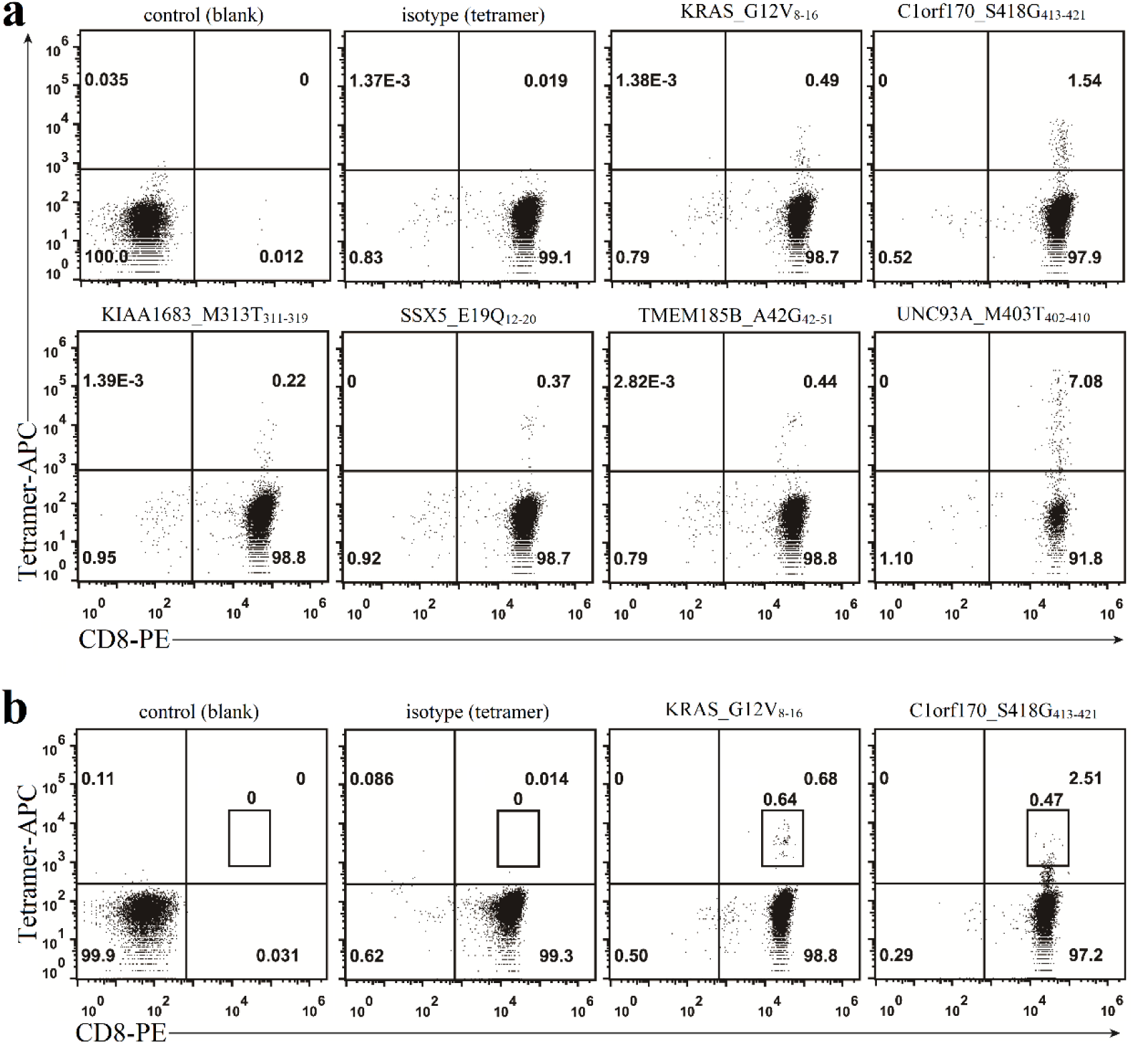
The proportion of epitope-specific CTL. Top 6 immunogenic peptides of KRAS_G12V_8-16_, C1orf170_S418G_413-421_, KIAA1683_M313T_311-319_, SSX5_E19Q_12-20_, TMEM185B_A42G_42-51_ and UNC93A_M403T_402-410_ were used to prepare pMHC tetramer. CTL, which had been co-cultured with peptide-loaded mDCs and confirmed to respond to peptide re-stimulation, were stained with CD8-PE and tetramer-APC. The fluorescence signal from cells was analyzed by FACS (a), and the CD8^+^pMHC tetramer^+^ cells that respectively responded to KRAS_G12V_8-16_ and C1orf170_S418G_413-421_ were sorted for single-cell RNA sequencing (b).

For T-cell receptor repertoires recognizing HLA-A*11:01 presented C1orf170_S418G_413-421_, a total of 9,243 cells (exactly GEM) were finally estimated from 17,800 sorted cells, and 14,042 TCR α chain and 9,346 TCR β chain amino acid sequences were respectively obtained, of which 8,826 pairs of TCR were produced. TCRα repertoire was preferentially biased toward usage of TRAV29DV5 gene (55.38%) and TRAV35 gene (37.35%) (Figure 6(a)), which corresponding mainly rearranged with TRAJ54 (99.88%) and TRAJ29 (Figure 6(b), and data not shown). The length distribution of CDR3α was preferentially restricted to 12 mer (55.64%) and 15 mer (39.09%) (Figure 6(c)), and the motif of 12-mer and 15-mer CDR3α was respectively highly conserved as CAASGGAQKLVF (Figure 6(g)) and CAGLLYNSGNTPLVF (Figure 6(h)), which was consistent with that of TRAV29DV5-TRAJ54 and TRAV35-TRAJ29. The gene usage of TCRβ repertoire was highly biased TRBV27 gene (87.47%) (Figure 6(d)), which mainly rearranged with TRBJ1-5 (99.91%) (Figure 6(e)). The length distribution of CDR3β was highly restricted to 15 mer (88.04%) (Figure 6(f)), and the motif of 15-mer CDR3β was highly conserved as CASSRDRGSNQPQHF (Figure 6(i)), which was consistent with that of TRBV27-TRBJ1-5. There were 298 diversities in the 8,826 TCRα/TCRβ pairs, but two clones accounted for a large proportion in the repertoire. 53.2% (4,694/8,826) T cells (named as Clonotype1) expressed TRAV29DV5-TRAJ54 and TRAV35-TRAJ29 containing TCRα and TRBV27-TRBJ1-5 containing TCRβ, and 31.6% (2,786/8,826) T cells (named as Clonotype2) expressed TRAV29DV5-TRAJ54 containing TCRα and TRBV27-TRBJ1-5 containing TCRβ (Table 3).

**Figure 6.**
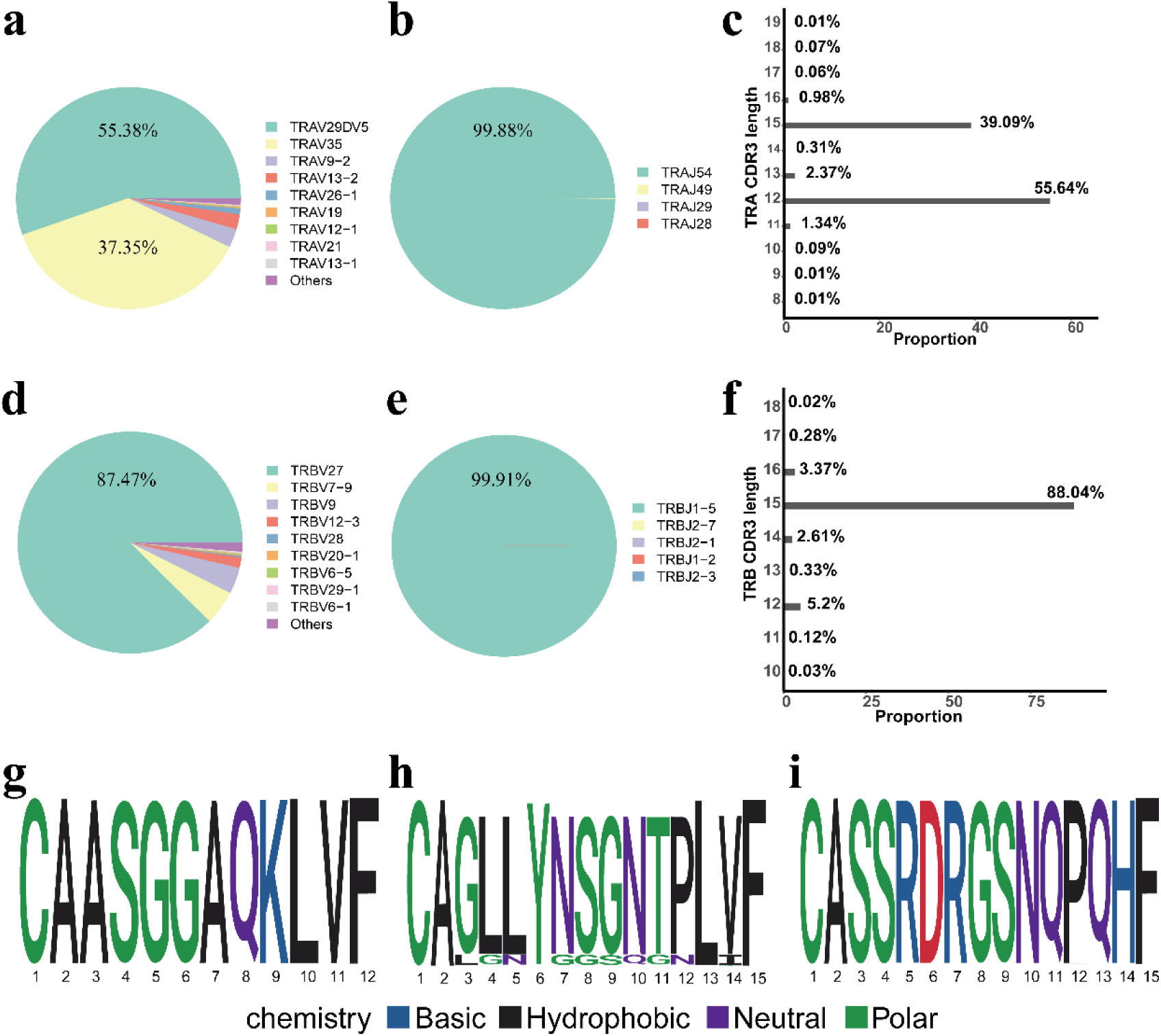
The characteristics of TCR repertoire recognized HLA-A*11:01 restricted C1orf170_S418G_413-421_. The frequency of usage gene of TRAV (a) and TRBV (d). The pie charts showed the top 9 usage genes, and the remaining usage genes were marked as “others”. The frequency of all the usage gene of TRAJ rearranged with TRAV29DV5 (b) and TRBJ rearranged with TRBV27 (e). The frequency of length distribution of CDR3α (c) and CDR3β (f). The sequence motif of 12-mer CDR3α (g), 15-mer CDR3α (h) and 15-mer CDR3β (i).

**Table 3.**
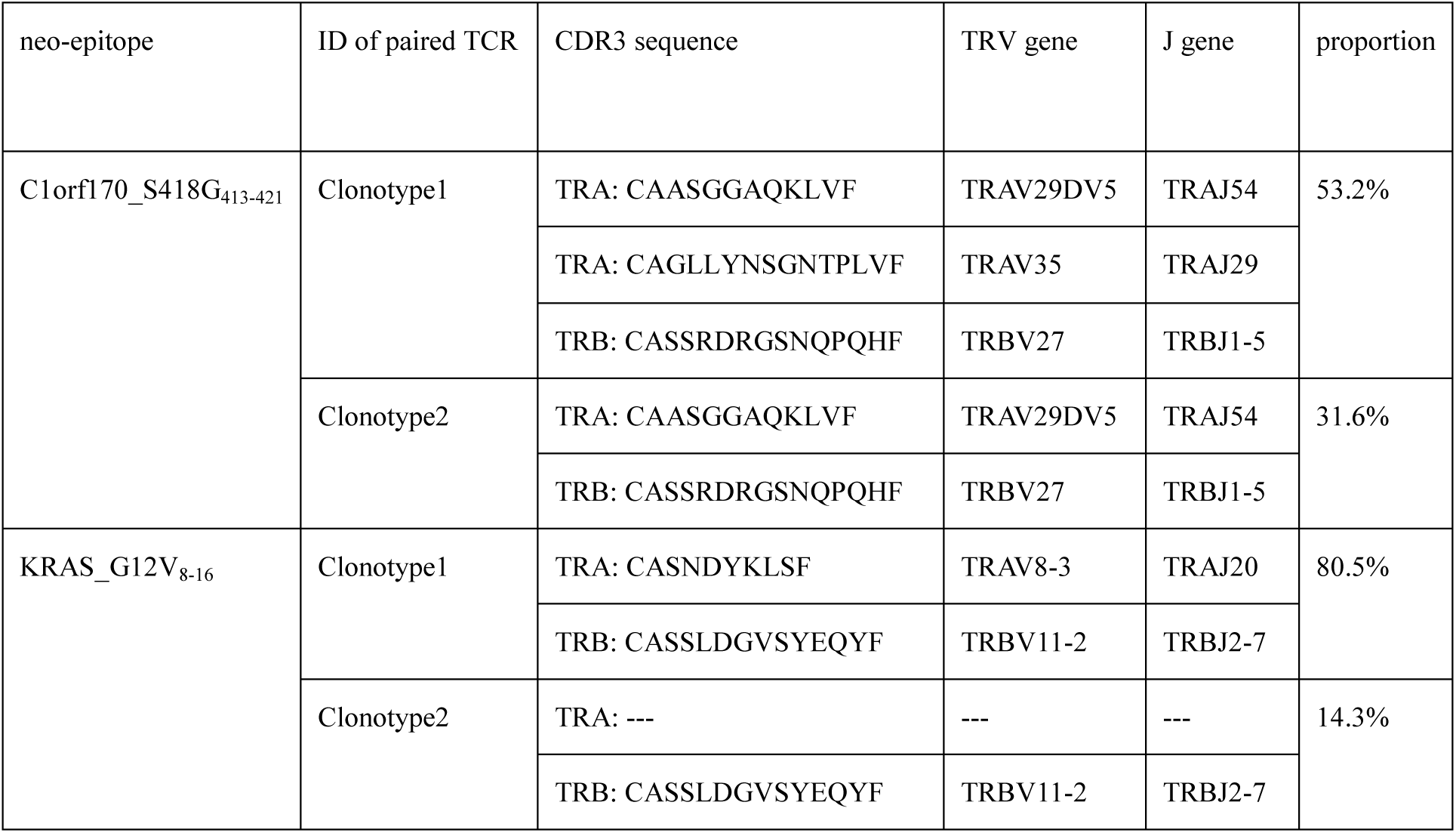
Top 2 TCR α and β chain pairs from two individual TCR repertoire

For T-cell receptor repertoires recognizing HLA-A*11:01 presented KRAS_G12V_8-16_, a total of 12,530 cells (exactly GEM) were finally estimated from 21,000 sorted cells, and 11,137 TCR α chain and 13,126 TCR β chain amino acid sequences were respectively obtained, of which 10,559 pairs of TCR were produced. TCRα and β repertoires respectively used immunodominant TRAV8-3 gene (81.69%) (Figure 7(a)), which mainly rearranged with TRAJ20 (99.69%) (Figure 7(b)), and TRBV11-2 gene (83.35%) (Figure 7(d)), which mainly rearranged with TRBJ2-7 (99.81%) (Figure 7(e)). The length distribution of CDR3α and CDR3β was respectively highly restricted to 10 mer (81.88%) (Figure 7(c)) and 14 mer (86.4%) (Figure 7(f)). The motif of 10-mer CDR3α and 14-mer CDR3β was respectively conserved as CASNDYKLSF (Figure 7(g)), which was consistent with that of TRAV8-3-TRAJ20, and CASSLDGVSYEQYF (Figure 7(h)), which was consistent with that of TRBV11-2-TRBJ2-7. There were 1,610 diversities in the 10,559 TCRα/TCRβ pairs, but two clones accounted for a large proportion in the repertoire. 80.5% (8,499/10,559) T cells (named as Clonotype1) expressed TRAV8-3-TRAJ20 containing TCRα and TRBV11-2-TRBJ2-7 containing TCRβ, and 14.3% (1,515/10,559) T cells (named as Clonotype2) expressed TRBV11-2-TRBJ2-7 containing TCRβ but were not detected TCRα (Table 3). Our results revealed that the dominant clone represented the usage genes and the CDR3 motifs of the TCR repertoire.

**Figure 7.**
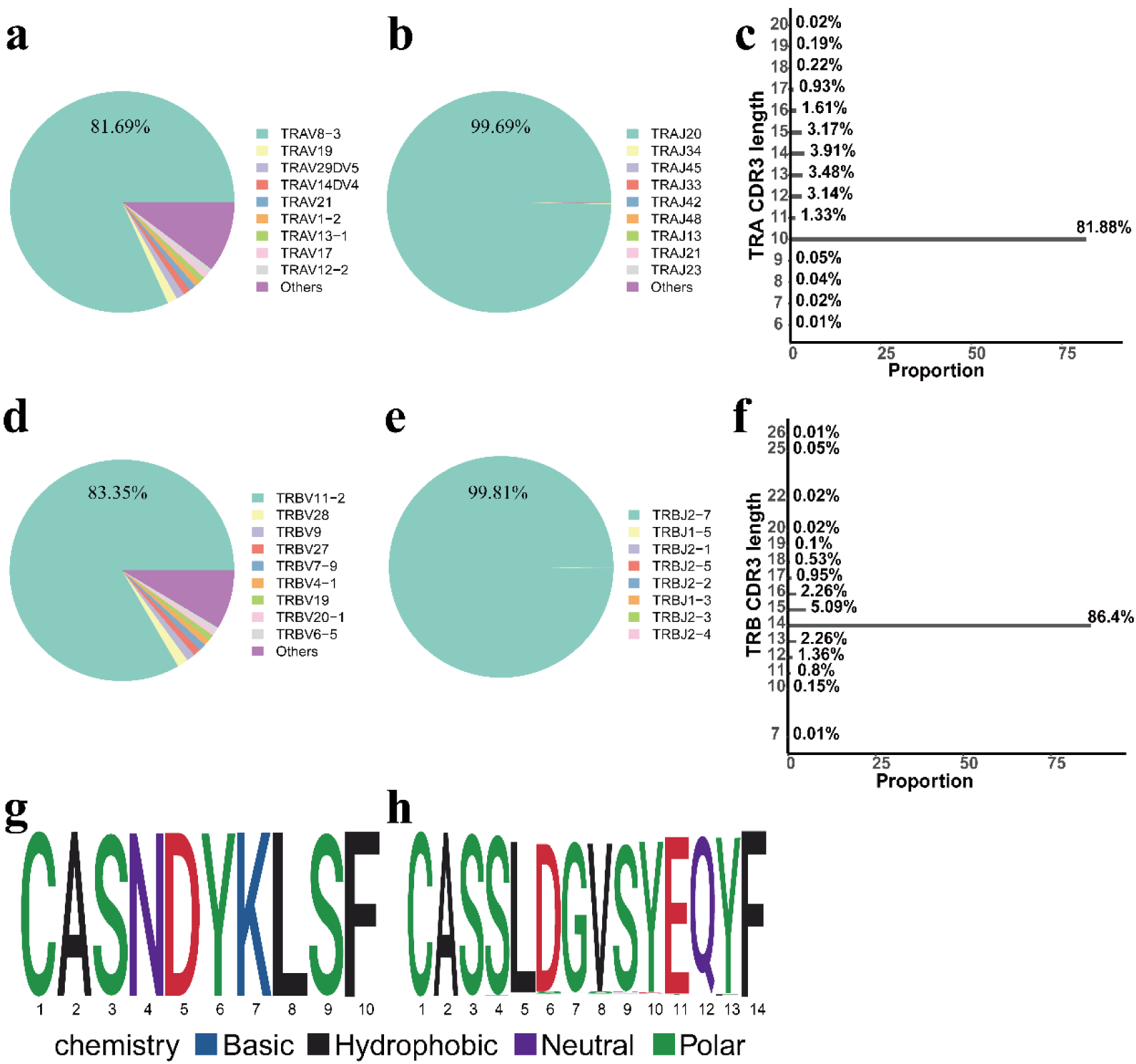
The characteristics of TCR repertoire recognized HLA-A*11:01 restricted KRAS_G12V_8-16_. The frequency of usage gene of TRAV (a) and TRBV (d). The pie charts showed the top 9 usage genes, and the remaining usage genes were marked as “others”. The frequency of all the usage gene of TRAJ rearranged with TRAV8-3 (b) and TRBJ rearranged with TRBV11-2 (e). The frequency of length distribution of CDR3α (c) and CDR3β (f). The sequence motif of 10-mer CDR3α (g) and 14-mer CDR3β (h).

The data that support the findings of this study have been deposited in the CNSA (https://db.cngb.org/cnsa/) of CNGBdb with accession code CNP0000518.

## Discussion

Cancer immunotherapy emerges as a very promising therapeutic approach for tumors. Different from chemotherapy, radiotherapy and targeted therapy, which directly target tumors, targeting the immune system offers the potential for durable activity and long-lasting survival outcomes [31]. It is widely accepted that anti-tumor immunity is especially mediated by the responses of tumor-specific T cells, which can effectively delete the primary tumor lesions and protest against metastases [31, 32]. T cells utilize TCR to recognize the short-peptide antigens bound in the groove of MHC molecules, and discriminate self and non-self, in which the short and cytosol-derived peptides mainly determine the specificity of the T cell-dependent immune response [33, 34].

During their carcinogenesis and progression, tumors usually obtain numerous somatic mutations. Mutant genes are translated into proteins and presented on cell surface by MHC, which result in arising neoantigens [11]. Neoantigens are uniquely produced by tumor cells, not totally found in normal tissues, and unparalleled tumor biomarkers [11]. Based on T cells recognizing neoantigens are not subject to thymic selection and central tolerance, neoantigen specific T cell with high-avidity is very likely to exist in the human body [8, 11]. Whole exome sequencing and RNA sequencing combined with bioinformatic pipelines make the reality of disclosing tumor-specific alterations with single nucleotide resolution and predicting neoantigens for cancer immunotherapy [35]. After pinpointing missense mutations and gene expression levels, peptides are assessed using various algorithms to predict binding affinity to MHC or presentation on MHC [20]. However, the vast majority of predicted neoantigens fail to turn up in tumors, and a handful is found to elicit T-cell responses [36]. Using five kinds of cancer patient-derived PBMCs, Chizu et al. only identified one immunogenic peptide from 26 mutant epitopes, which were predicted by NetMHC4.0 algorithm to have strong-binding capacity to HLA-A*24:02 [9]. Using four kinds of healthy donor-derived T cells, Stronen et al. showed T-cell reactivity toward 3-5 of 20 neoantigens, which were predicted by NetMHC3.2 or netMHCpan2.0 algorithm to have high predicted binding to HLA-A*02:01[37]. In a small group of patients with stages III and IV melanoma, Ott et al. demonstrated only 16% neoantigens, which were predicted by NetMHCpan-2.4 algorithm, were recognized by CD8^+^ T cells [38]. Zhang et al. found a significant T-cell response in two of nine neoantigens for one breast cancer patient, and one of eight neoantigens for the other two patients with breast cancer, in which neoantigens were predicted by NetMHC-3.2 algorithm [39]. Therefore, we considered that none of the current algorithms was perfect, and it was necessary to simultaneously use multiple algorithms to increase the accuracy of peptide binding affinity prediction. In the present study, we respectively used peptide-MHC binding-affinity prediction algorithms, including NetMHC-4.0, NetMHCpan-3.0, NetMHCpan-4.0, PSSMHCpan-1.0, PickPocket-1.0 and SMM, and EPIC algorithm that predicted epitope presentation to evaluate the extracted peptides. Finally, we selected 25 candidate peptides, which simultaneously met the conditions of the frequency of SNV being over 5 out of 321 patients and InDel being over 2 out of 321 patients, IC50 value being less than 50 nM by at least three software packages and EPIC score value being more than 0.9 (Table 1), to assay their characteristics of presentation and inducing cytotoxic T cells. We found that 11 out of 25 (44%) predicted epitopes were proved to be presented by HLA-A*11:01 allele through MS (Figure 2), and 11 out of 25 (44%) predicted epitopes induced specific CTL to secrete IFN-γ through ELISPOT assay (Figure 4), and 20 out of 25 (80%) predicted epitopes could either be presented or have immunogenicity (Figure 2 and Figure 4). However, it was a pity that, except the positive epitope (KRAS_G12V_8-16_), only 2 out of 25 (8%) predicted epitopes were analyzed not only to be endogenously expressed in tumor cells, but also to induce a T-cell response (Figure 2 and Figure 4).

At present, MS-based approach is the relatively unbiased methodology to identify the repertoire of peptides, which are naturally processed and presented by MHC molecules *in vivo*, from human cancer cell lines, tumors and healthy tissues and body fluids [40]. Although the use of MS-based immunopeptidomics would reduce the false positive number of predicted *in silico* neoantigens, and ensure highly accurate and reliable assignment of neoantigen’s sequences, neoantigens have not been regularly and sensitively disclosed by MS compared with TAAs [35, 40]. Based on the limited sensitivity, the false negative neoantigens that are naturally presented but not detected by MS are expected for immunopeptidomics. Due to the reports that relatively large biological samples, abundance of proteins containing specific sequences, expression in mono-allelic cells, proteasomal processing and the MITD trafficking signal for siting in endolysosomal compartments are important facts for discovering the presented peptides [22, 25, 40, 41], Five tandem minigenes, which each encoded six neo-antigenic peptides, was operated by CMV promoter and linked with the MITD trafficking signal at the C terminal (Figure 1(a)), were constructed and transfected into HLA-A*11:01 mono-allelic K562 cells. We found that over 85% HLA-A*11:01 mono-allelic K562 cells highly expressed the tandem minigenes (Figure 1(b)), and used 1×10^9^ cells to extract and purify the peptides, which were followed by analysis with a mass spectrometer. Finally, except the positive epitope (KRAS_G12V_8-16_), we confirmed that 11 out of 25 predicted neoantigens were naturally processed and presented by HLA-A*11:01 allele, in which the positive rate was up to 44% (11/25) (Figure 2). However, the remaining neo-peptides were not detected in our system. Recently, Muhammad Ali et al. proved that minimizing the formation of irrelevant immunogenic peptides could increase the targeted epitopes to bind HLA, and the order of arranged epitopes in the tandem minigene was involved in the efficiency of antigen presentation [27]. In our present study, the predicted neoantigens in the tandem minigene each had 27 amino acids with the mutation at position 14 (Table 2), which may result in the formation of other high-affinity irrelevant and competitive presented peptides, and were randomly combined in the tandem minigene. Therefore, we considered that the proper length and the appropriate order of desired neoantigens in the tandem minigene may improve the probability of the remaining neo-peptides being detected by MS.

The immunogenicity of predicted neoantigens is usually detected through the response of TIL [11, 12]. Moreover, although the frequency of the neoantigen-reactive T cells in the peripheral blood is quiet low compared with that in the tumor sample, neoantigen-specific T cells, which derive from circulating CD8^+^ memory T cells of cancer patients or circulating CD8^+^ naïve T cells of healthy donors, can be enriched after being *in vitro* co-cultured with the cognate DCs pre-loaded neoantigens, and their response to recognize the corresponding neoantigens can be detected by conventional experimental methods [12, 27, 42]. Therefore, we generally adopted one of the proven methods to detect the immunogenicity of our predicted neoantigens, in which the circulating bulk CD8^+^ T cells from one healthy donor were stimulated for two rounds by cognate DCs pre-loaded neoantigens. Except the positive epitope (KRAS_G12V_8-16_), we found 11 out of 25 (44%) predicted neoantigens that were pre-loaded on T2 cells could induce co-cultured CTL to secrete IFN-γ (Figure 4), and the frequencies of the specific CTL, that produced over 10 folds IFN-γ spots than the negative control, were estimated to range between 0.22% and 7.08% (Figure 5(a)). However, most of the presented neoantigens were not detected to be immunogenic in our study. As approximately only one in 10^5^-10^6^ T cells is specific for a given antigen in the lymph node, and circulating CD8^+^ memory T cells from healthy donors do not contribute to the production of neoantigen-responsive T cells, to enrich circulating CD8^+^ naïve T cells from healthy donors before priming can enhance the probability of specific T cells encountering DCs presenting the cognate neoantigen [27, 43]. Based on the donor-dependent variability and sufficient diversity of the human TCR repertoire, several donors need to be screened for identifying the immunogenicity of a novel candidate neoantigen [27]. Therefore, we considered that CTL, which were expanded from CD8^+^ naïve T cells of other healthy donors, may respond to the remaining immunogenic negative neo-peptides.

In conclusion, except the positive epitope (KRAS_G12V_8-16_), our results revealed several common actual T-cell neo-epitopes of CRC, which would be developed as the universal targets for CRC immunotherapy in the form of vaccines based on peptide, RNA, DNA and DCs and therapies based on adoptive TCR transgenic T cells.

## Abbreviations

CRC: colorectal cancer
MMR: deficient mismatch repair
MSI-H: highly microsatellite instable
pMMR: proficient mismatch repair
MSS: microsatellite stable
ADC: autologous tumor lysate DC
CAR: chimeric antigen receptor
TAA: tumor associated antigen
CEA: carcinoembryonic antigen
HER2: human epidermal growth factor receptor-2
TSA: tumor-specific antigen
MHC: major histocompatibility complex
TMB: tumor mutational burden
TCR: T-cell receptor
PRM: parallel reaction monitoring
TIL: tumor infiltrating lymphocyte
MS: mass spectrometry
ATCC: American Type Culture Collection
PBMC: peripheral blood mononuclear cell
ICGC: International Cancer Genome Consortium
COCA-CN: China-Colorectal Cancer Project
InDel: insertions or deletion
EPIC: Epitope Presentation Integrated prediction
MITD: MHC class I trafficking signal
CTL: cytotoxic lymphocyte
ELISPOT: enzyme-linked immunospot
FACS: fluorescence-activated cell sorting
GEM: Gel Bead in Emulsion

## Acknowledgments

We thank Weipeng Hu from BGI-GenoImmune for technical assistance on this study.

